# Broadband aperiodic components of local field potentials reflect inherent differences between cortical and subcortical activity

**DOI:** 10.1101/2023.02.08.527719

**Authors:** Alan Bush, Jasmine Zou, Witold J. Lipski, Vasileios Kokkinos, R. Mark Richardson

## Abstract

Information flow in brain networks is reflected in intracerebral local field potential (LFP) measurements that have both periodic and aperiodic components. The 1/f^χ^ broadband aperiodic component of the power spectra has been shown to track arousal level and to correlate with other physiological and pathophysiological states, with consistent patterns across cortical regions. Previous studies have focused almost exclusively on cortical neurophysiology. Here we explored the aperiodic activity of subcortical nuclei from the human thalamus and basal ganglia, in relation to simultaneously recorded cortical activity. We elaborated on the FOOOF (fitting of one over f) method by creating a new parameterization of the aperiodic component with independent and more easily interpretable parameters, which allows seamlessly fitting spectra with and without an *aperiodic knee*, a component of the signal that reflects the dominant timescale of aperiodic fluctuations. First, we found that the aperiodic exponent from sensorimotor cortex in Parkinson’s disease (PD) patients correlated with disease severity. Second, although the aperiodic knee frequency changed across cortical regions as previously reported, no aperiodic knee was detected from subcortical regions across movement disorders patients, including the ventral thalamus (VIM), globus pallidus internus (GPi) and subthalamic nucleus (STN). All subcortical region studied exhibited a relatively low aperiodic exponent (χ^STN^=1.3±0.2, χ^VIM^=1.4±0.1, χ^GPi^=1.4±0.1) that differed markedly from cortical values (χ^Cortex^=3.2±0.4, f_k^Cortex^_=17±5 Hz). These differences were replicated in a second dataset from epilepsy patients undergoing intracranial monitoring that included thalamic recordings. The consistently lower aperiodic exponent and lack of an aperiodic knee from all subcortical recordings may reflect cytoarchitectonic and/or functional differences between subcortical nuclei and the cortex.

**Significance Statement:** The broadband aperiodic component of local field potentials is a useful and reproducible index of neural activity. Here we refined a widely used phenomenological model for extracting aperiodic parameters, with which we fit cortical, basal ganglia and thalamic intracranial local field potentials, recorded from unique cohorts of movement disorders and epilepsy patients. We found that the aperiodic exponent in motor cortex is higher in Parkinson’s disease patients with more severe motor symptoms, suggesting that aperiodic features may have potential as electrophysiological biomarkers for movement disorders symptoms. Remarkably, we found conspicuous differences in the broadband aperiodic components of basal ganglia and thalamic signals compared to those from neocortex, suggesting that the aperiodic neural timescale of subcortical LFPs is slower than that in cortex.

## Introduction

From the inception of EEG, understanding the neurophysiology of the oscillatory electrical activity—periodic activity of defined frequencies which is sustained for more than one period—has been a paramount goal (Berger, 1929). These neural oscillations have been found to be widespread, spanning all brain regions and frequency bands, and correlate with many aspects of brain function and dysfunction (Basar and Güntekin, 2013; Engel et al., 2001). The study of neural oscillations has also been facilitated by commonly used methods like Fourier or wavelet transforms, which can decompose any signal into a sum of oscillatory components. However, the existence and mathematical validity of these decompositions does not imply that all brain activity arises from neural oscillations. Indeed, processes that create fluctuations in the signal with no underlying oscillatory component give rise to characteristic power spectra when analyzed by these same methods.

Local field potentials (LFPs) reflect the ensemble activity of ionic currents of populations of cells in the vicinity of the electrode (Lindén et al., 2010; Nunez and Srinivasan, 2006). The most salient feature of the frequency power spectral density (PSD) of LFPs is the decline of power with frequency, a feature termed the 1-over-f (1/f^χ^) “background noise” of the spectra. Studies using LFPs commonly remove the 1/f^χ^ broadband component by normalization and focus on modulation of power at specific frequency bands. To contrast the periodic nature of neural oscillations, the 1-over-f component is referred to as *broadband aperiodic activity*.

Until recently, aperiodic activity has been largely ignored or regarded as noise, perhaps due to inadequate computational tools and theoretical framework. In pioneering work, Miller et al. fitted a parametric description of the broadband aperiodic component to human electrocorticography (ECoG) PSD (Miller et al., 2009). The extraction of the aperiodic exponent χ has been greatly facilitated by the development of methods like the irregular-resampling auto-spectral analysis (IRASA) (Wen and Liu, 2016) and fitting of one-over-f (FOOOF) (Donoghue et al., 2020; Haller et al., 2018). The latter fits the periodic component of the spectrum as a superpositions of gaussians and parameterizes the aperiodic component as *P_aper_* = *A*/(*k* + *f^χ^*), with an offset **A**, an aperiodic exponent **χ**, and an optional knee parameter **k** (Donoghue et al., 2020; Haller et al., 2018) (see also Supplementary Materials). Note that this method requires an *a priori* decision of whether to use the knee parameter or not.

Using these methods, a recent body of work explored correlations of aperiodic parameters with different behavioral, physiological, and pathophysiological states, and anatomical regions. The cortical aperiodic exponent **χ** decreases with age (Dave et al., 2018; Voytek et al., 2015), increases under anesthesia and during sleep (Colombo et al., 2019; Lendner et al., 2020; Miskovic et al., 2019; Muthukumaraswamy and Liley, 2018), and differs across cortical regions (Chaoul and Siegel, 2021; Muthukumaraswamy and Liley, 2018). Likewise, the knee **k** of the spectra (i.e., the frequency at which the 1/f^χ^ decline of power with frequency begins) also has a spatial structure in the cortex (Gao et al., 2020a). Thus, aperiodic parameters are useful population-average measures of neural activity.

Given the importance of understanding the cortical-subcortical neural dynamics that underly normal human behavior and symptoms of brain diseases, we explored differences in the parameters of the aperiodic component of LFPs recorded from unique cohorts of neurosurgical patients. We elaborated on the parameterization of the broadband aperiodic component developed by (Donoghue et al., 2020; Haller et al., 2018) to obtain a model with better defined aperiodic parameters that avoids *a priori* assumptions on the presence of an aperiodic knee. We used this model to explore (across patients) the relation of cortical aperiodic activity with movement disorder pathophysiology and cortical anatomy, in movement disorders patients undergoing deep brain stimulation (DBS) surgery. We then performed within subject comparisons of aperiodic parameters in thalamic and basal ganglia nuclei to those in cortex, including a second cohort of patients with drug-resistant epilepsy undergoing intracranial monitoring.

## Methods

### Participants

Movement disorder patients undergoing intracranial electrode implantation for deep brain stimulation therapy participated in a speech production task (Bush et al., 2021), for which the baseline periods were analyzed in this study. One or two high-density subdural electrocorticography (ECoG) strips were temporary placed through the standard burr hole, targeting the left superior temporal gyrus (covering also the ventral sensorimotor cortex) and left inferior frontal gyrus. ECoG electrodes were removed at the end of the surgery. Dopaminergic medication was withdrawn the night before surgery. All procedures were approved by the University of Pittsburgh Institutional Review Board (IRB Protocol #PRO13110420) and all patients provided informed consent to participate in the study. The following cohorts of movement disorder patients participated in the study: 29 Parkinson’s disease patients (21M/8F, 65.6±7.1 years) undergoing awake subthalamic (STN) DBS surgery, all of which had ECoG recordings and 14 of which had simultaneous ECoG and DBS lead recordings; 5 Parkinson’s disease patients (5M/0F, 69.1±5.7 years) undergoing awake pallidal (GPi) DBS surgery, of which 4 had ECoG recordings and 3 had simultaneous ECoG and DBS lead recordings; 22 essential tremor patients (11M/11F, 65.3±9.7 years) undergoing awake thalamic (Vim) DBS surgery, of which 20 had ECoG recordings and 11 had simultaneous ECoG and DBS lead recordings.

Additionally, we analyzed awake restfulness data from 8 epilepsy patients (5M/3F, age: 18±11 years) undergoing stereo-EEG (sEEG) intracranial monitoring for epilepsy with additional electrodes implanted in the thalamus. This study was approved by the Massachusetts General Hospital (Boston, MA) Institutional Review Board (IRB Protocol #2020P000281).

### Neural recordings

Figure 1 and Table S1 describe the electrodes used in this study. Signals from ECoG electrodes and DBS leads were acquired at 30kHz (filtered between 1 Hz and 7.5 kHz) with a Grapevine Neural Interface Processor equipped with Micro2 Front Ends (Ripple LLC, Salt Lake City, UT, USA). ECoG and DBS lead recordings were referenced to a subdermal scalp needle electrode positioned approximately on Cz. The sEEG signals were recorded at 1 kHz sampling rate using a 128-channel Xltek digital video-EEG system (Natus Medical Incorporated, Pleasanton, CA). sEEG recordings were referenced to an EEG electrode placed extracranially (C2 vertebra or Cz).

**Figure 1.**
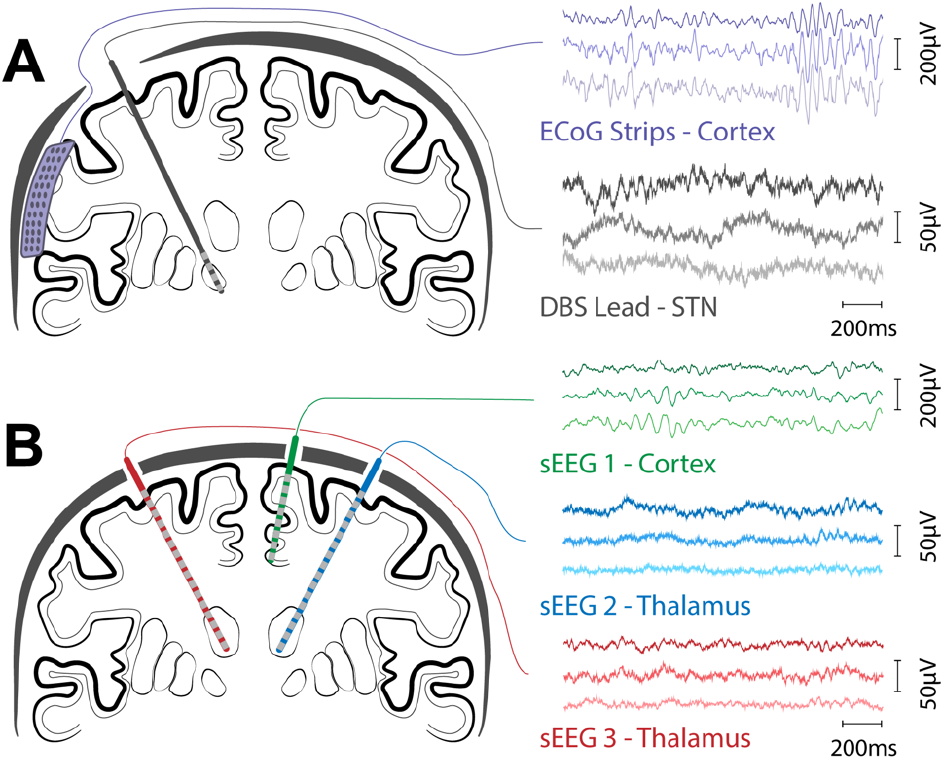
Schematic representation of coronal view of electrode montages. A) Movement disorder patients undergoing DBS implantation surgery with simultaneous multichannel recordings from DBS leads and ECoG strips. B) Epilepsy patients undergoing intracranial monitoring with multichannel sEEG electrodes, some of which target thalamic nuclei.

### Electrode localization

DBS electrodes were localized using the Lead-DBS localization pipeline (Horn et al., 2019). Briefly, a pre-operative anatomical T1 weighted MRI scan was co-registered with a post-operative CT scan. Position of individual contacts were manually identified based on the CT artifact and constrained by the geometry of the DBS lead used. This process rendered the coordinates for the leads in each subject’s native space. The position of the ECoG strips were calculated from intra-operative fluoroscopy as described in (Randazzo et al., 2016). Briefly, the cortical surface was reconstructed from the pre-operative MRI using FreeSurfer (Fischl et al., 2002) and a model of the skull and stereotactic frame was reconstructed from the intra-operative CT scan using OsiriX (osirix-viewer.com). The position of the frame’s tips on the skull and the implanted DBS leads were used as fiducial markers. The models of the pial surface, skull and fiducial markers were co-registered, manually rotated and scaled to align with the projection observed in the fluoroscopy. Once aligned, the position of the electrodes in the ECoG strip were manually marked on the fluoroscopy image and the projection of those position to the convex hull of the cortical surface was defined as the electrode location in native space. The coordinates were then regularized based on the known layout of the contacts in the ECoG strip (github.com/Brain-Modulation-Lab/ECoG_localization). All coordinates were then transformed the ICBM MNI152 Non-Linear Asymmetric 2009b space (Fonov et al., 2011) using the Symmetric Diffeomorphism algorithm implemented in the Advanced Normalization Tools (Avants et al., 2008).

Epilepsy patients were implanted with commercially available 8 – 16 contact electrodes (PMT Corporation, MN, USA; AdTech Medical Instrument Corporation, WI, USA). Electrode trajectories were tailored for each patient according to the surgical hypothesis and contact locations were determined by either post-implantation MRI, co-registration of the pre-operative T1 MRI with the post-implantation CT using Brainstorm (Tadel et al., 2011).

Anatomical labels were assigned to each electrode based on the HCP-MMP1 atlas (Glasser et al., 2016) for cortical electrodes, and the Morel (Morel, 2007) and DISTAL (Ewert et al., 2018) atlases for subcortical electrodes.

### Electrophysiological data preprocessing and power spectrum estimation

Data recorded during DBS surgeries was processed using custom code based on the FieldTrip (Oostenveld et al., 2011) toolbox implemented in MATLAB, available at (github.com/Brain-Modulation-Lab/bml). Data was low pass filtered at 250Hz using a 4^th^ order non-causal Butterworth filter, down-sampled to 1 kHz and stored as continuous recordings in FieldTrip datatype-raw. No notch filter was applied. Electrodes were common average referenced per head-stage connector and electrode type. Power spectral density (PSD) was estimated using the Welch method (Welch, 1967), using 1 s time windows over the inter-trial baseline periods of the speech task with a 500ms overlap. The median PSD across all baseline periods was calculated for subsequent analysis. sEEG data recorded for epilepsy monitoring, was processed using the MNE toolbox in python. Recordings were bipolar referenced and PSDs were estimated using the Welch method by calculating periodograms for a sliding window of two seconds and overlap of 100 ms.

### Spectral parameterization

We elaborated upon the spectral parameterization introduced by (Donoghue et al., 2020; Haller et al., 2018) to capture the frequency domain characteristics of electrophysiological data. This parameterization decomposes the log-power spectra **log(P(*f*))** into a broadband aperiodic component **log(L(*f*))** and the summation of **N** narrowband periodic components which are each modelled as a Gaussian.

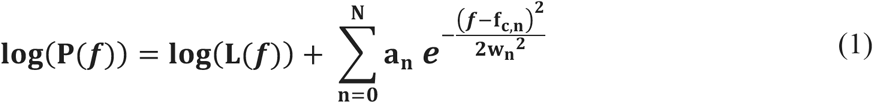

where *f* is the frequency, **a_n_** is the power, **f_c,n_** the center frequency and **w_n_** is the width of the Gaussian **n** (i.e., the standard deviation). Gaussians were used to model physiological oscillations and spectral artifacts like line noise. This approach was preferred over using notch filters as spectra with notches were not adequately fitted by the proposed model. In this work we propose a new parameterization of the aperiodic component defined as

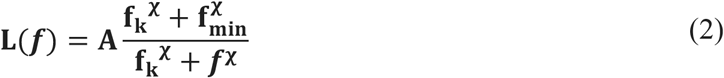

where **A** is the broadband offset and can be interpreted as the power fitted at the minimal frequency of interest **f_min_**, that is, the smallest positive frequency for which power can be reliably estimated based on acquisition, preprocessing, and PSD estimation method and parameters. For the current work, it was determined as **f_min_ = max{f**_HP_**, f**_s_**/m}**, the largest between **f**_HP_, the cutoff frequency of the high-pass filter applied at acquisition (or preprocessing), and the smallest positive frequency calculated by the Welch method **f_s_**/**m**, where **f_s_** is the sampling rate and **m** the number of samples in the Welch window. The parameter **f**_k_ is the knee frequency which (for **f**_k_ ≫ **f_min_**) can be interpreted as the frequency at which the power decays to approximately A/2. The rate at which the power decreases for frequencies above **f**_k_ is defined by the aperiodic slope **χ**. We also modified the original algorithm proposed by (Haller et al., 2018) to scan **f_*k*_** logarithmically, therefore ensuring positive values. This change also allows the full model to adequately fit cases in which there is no knee in the PSD by converging to **f**_*k*_ ≪ f_*min*_. For computational reasons we restricted the range of **f_*k*_** from **f_*min*_/10** to **f_*Nyquist*_**. See the supplementary materials for a discussion on the advantages of using this parameterization over the original one proposed by (Donoghue et al., 2020; Haller et al., 2018).

Additionally, we modified the cost function (J) of the fitting procedure by adding to the mean squared error term a regularization term that penalizes the integral of the gaussians over negative frequencies (Equation 3),

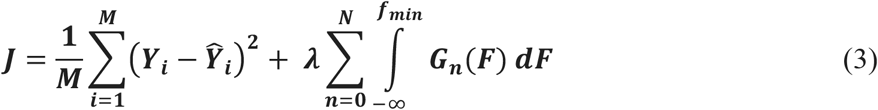

where ***Y*_i_** is the log-power estimated by the Welch method at frequency **f_i_**, and *Ŷ*_i_ is the value fitted by the model. The second term was added to prevent Gaussian peaks to extend beyond the fitting range, which can affect the estimation of aperiodic component (Gerster et al., 2022). The regularization parameter *λ* was empirically adjusted for each dataset. Algorithm development and analyses for this work were done in Python. Scripts and packages are available at github.com/Brain-Modulation-Lab/fooof/tree/lorentzian.

### Statistical analysis

We performed statistical analyses in R. Base functions were used for correlation tests, paired t-tests, linear models, and Fisher exact test for count data. The *coin* package was used for permutation tests (Hothorn et al., 2008), *lmerTest* for linear mixed effects models (Kuznetsova et al., 2017) and *multcomp* for multiple comparisons (Bretz et al., 2011).

## Results

To explore differences between the aperiodic components of cortical and basal ganglia or thalamic LFPs, we elaborated upon the FOOOF method (Donoghue et al., 2020; Haller et al., 2018) by incorporating a new Lorentzian-like parameterization of the broadband aperiodic component, changing the way parameters are scanned and adding a regularization term (see methods and supplementary materials for details). These changes result in more easily interpretable parameters, with well-defined units and better parameter identifiability (Cedersund and Roll, 2009) (Figure S1). These modifications also allow fitting of the same model to power spectra with qualitatively different profiles. In the original description, parameterization required an *a priori* selection of one of two possible models (with or without a “knee” parameter); our modifications allow seamless fitting of either case with the same model.

First, to assess the performance of the novel parameterization, we fitted the power spectra of ECoG recordings acquired from movement disorder patients undergoing awake DBS implantation surgery. Baseline epochs recorded during rest periods in a speech production task were used for this analysis. The novel parameterization fits the data as well as the original implementation (Figure 2a); R^2^ values of both models are virtually identical and tightly cluster at values above 0.975 (Figure 2a inset). However, the aperiodic parameters for the novel formulation do not show the strong collinearity observed for the parameters of the original model (Figure 2b and S2). In this context, collinearity is indicative of poor parameter identifiability, leading to larger uncertainties of the parameters (Cedersund and Roll, 2009). (Note however that there is a residual correlation between the aperiodic knee and the exponent of the spectra, Figure S2b). Our novel formulation also better constrains the range of values of the parameters, for example the aperiodic offset spans 6 orders of magnitudes for the original model but only 2 in the novel formulation (Figure 2b).

**Figure 2.**
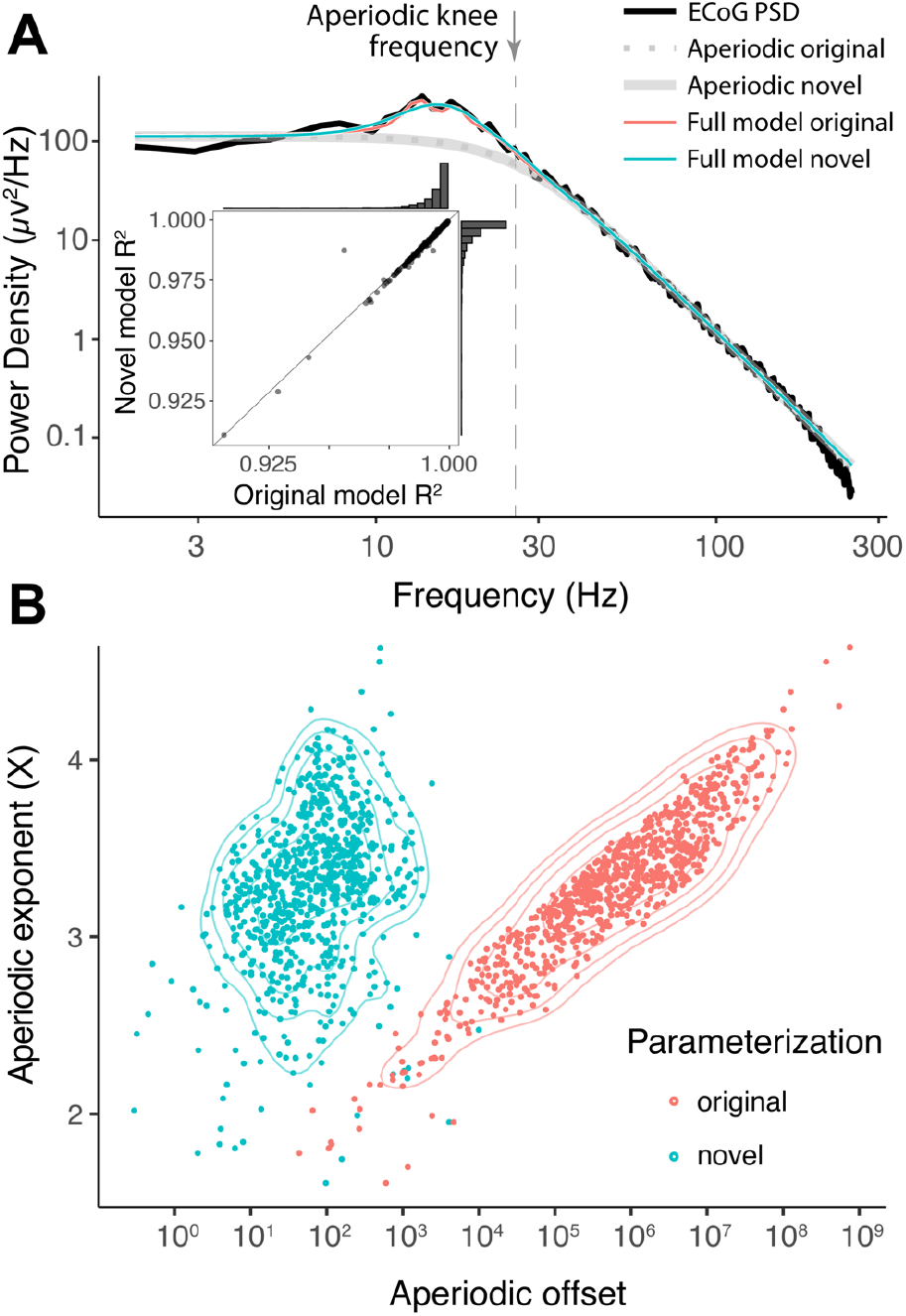
The novel parameterization of the aperiodic component avoids collinearity between parameters. A) Representative example of cortical power spectra with fits from original and novel models. The inset shows the correlation between R^2^ values for both models, and their univariate distribution in data from PD participants. B) Aperiodic exponent vs. offset parameters for ECoG recordings from PD patients, for the original (red) and novel (blue) parameterizations. Contour lines represent the 5%, 10%, 20%, 40% and 80% percentiles of 2D kernel density estimation.

Using this new parameterization, we explored cortical aperiodic activity from ECoG recordings in PD patients undergoing STN-DBS implantation. Across participants, electrodes covered the left inferior frontal cortex, precentral and postcentral gyrus, superior and middle temporal gyrus (blue dots in Figure 3a). Interestingly, we found a significant positive correlation between the pre-operative UPDRS-III ON score and the aperiodic exponent from Rolandic and premotor cortical areas (r=0.4, p=0.036, Pearson correlation, Figure 3b), but not from other cortical regions. However, no significant correlation was found with the UPDRS-III OFF score (r=0.25, p=0.22), nor the UPDRS-III ON/OFF percent change (r=0.33, p=0.13, Figure S3). There was no significant difference in cortical aperiodic parameters extracted from ECoG of PD and essential tremor (ET) patients for the exponent (p=0.28, permutation test), knee frequency (p=0.68) or offset (p=0.28, Figure 3c). Therefore, we pooled data across these two cohorts for subsequent analyses. There was no significant correlation of the aperiodic components with age (p=0.44, Pearson correlation). Note that the age range of this cohort (43-79 yrs) does not include the younger adult group (18-30 yrs) from previous studies (Dave et al., 2018; Voytek et al., 2015). We grouped electrodes according to the multimodal parcellation 1 atlas (HCP-MMP1, (Glasser et al., 2016), Figure 3d), and used a mixed effects model to account for subject-to-subject variability. In line with recent reports (Chaoul and Siegel, 2021; Gao et al., 2020a; Muthukumaraswamy and Liley, 2018) we found significant differences in aperiodic parameters across cortical regions (Figure 3e, Table S2). We observed that the aperiodic knee frequency in primary sensory cortex “1” was significantly greater than that observed in frontal regions “8Av” (p=0.014) and “6r” (p=0.017), opercula area 4 “OP4” (p<0.01) and secondary auditory cortex “A5” (p<0.01, Tukey’s Contrast for region effect).

**Figure 3.**
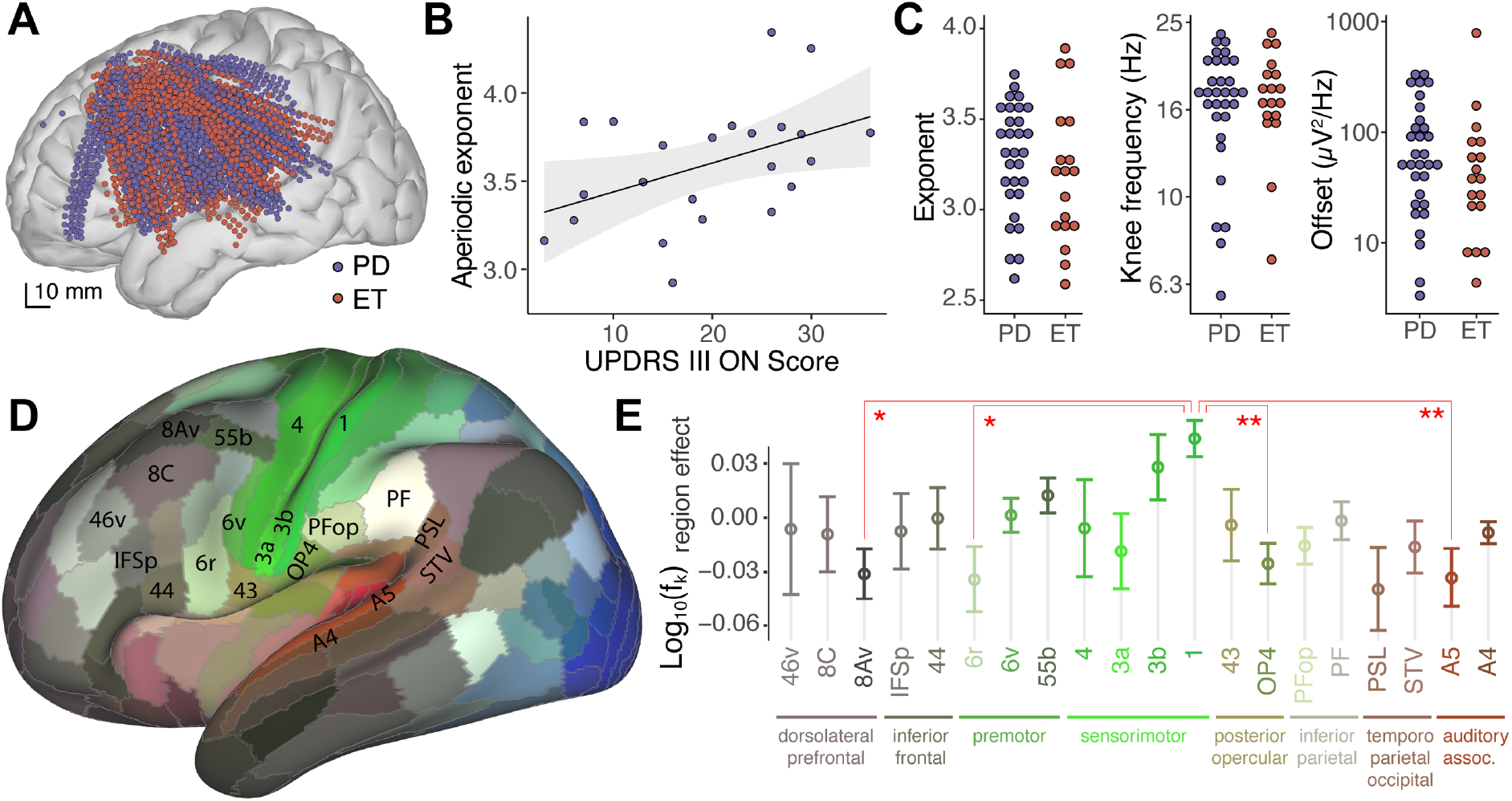
Cortical aperiodic parameters correlate with PD severity and anatomical regions. A) Anatomical localization of ECoG electrodes used to record cortical activity from Parkinson’s disease (PD, blue) and essential tremor (ET, red) patients undergoing DBS surgery. B) Median cortical aperiodic exponent from Rolandic and premotor ECoG recordings in PD patients undergoing STN-DBS surgery vs. preoperative UPDRS-III ON score. Shaded region represents CI_95_. C) Median values of cortical aperiodic exponent (left), knee frequency (center) and offset (right) for each subject, color coded by diagnosis. D) Lateral view of cortical parcellation defined by MMP1 on an inflated brain, colored according to fMRI response to visual (blue), auditory (red) or somatosensory (green) tasks (Glasser et al., 2016). E) Aperiodic knee frequency cortical region effect (after accounting for subject effect) vs. anatomical regions, as defined in the MMP1 atlas, for regions recorded by electrodes from 10 or more subjects. To avoid effects from differences in sampling density, statistics were done on the average per region per subject. Error bars indicate the SEM across subjects. Colors as in D.

Next, we explored the aperiodic potentials from subcortical recordings acquired through the DBS leads. For PD patients, DBS leads targeted the dorsal-posterior-lateral portion of the subthalamic nucleus (STN) or the inferior-posterior-lateral globus pallidus internus (GPi), whereas for ET patients leads targeted the ventral intermedius (VIM) nucleus of the thalamus (Figure 4a). In contrast to what was observed for cortical recordings, no obvious ‘knee’ was apparent in power spectra from the STN, VIM or GPi (Figure 4b, 4c and 4d); the aperiodic component of extracellular potentials for these subcortical structures decreases with frequency starting from the minimal frequency acquired. These qualitative differences with ECoG PSDs could be due to the different electrode types (see Table S1 for details), reflect underlying electrophysiology, or a combination of both effects (see discussion). To quantify these differences, we fit subcortical power spectra using the same model as for cortical data (Figure 4b, 4c and 4d).

**Figure 4.**
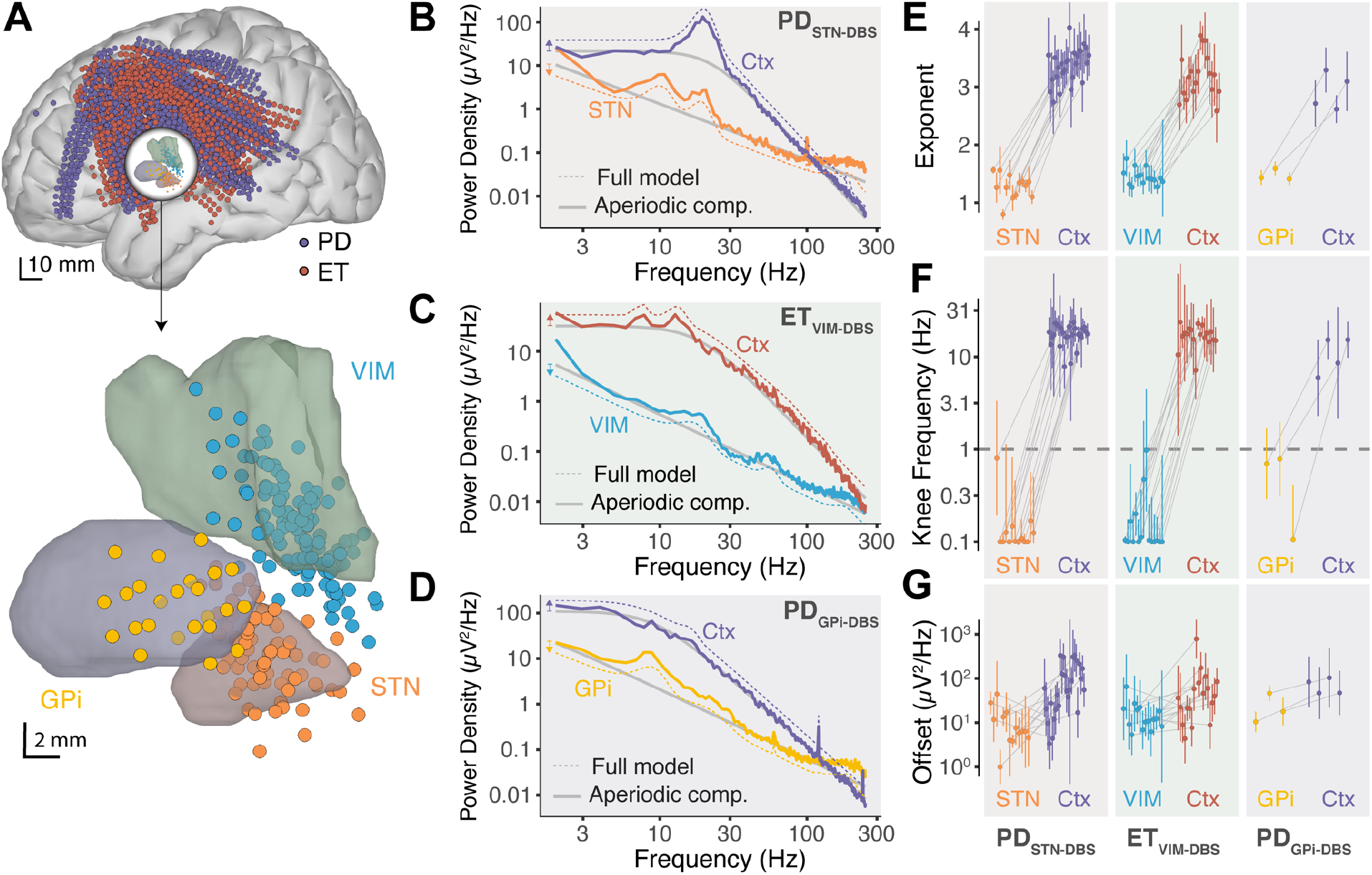
Power spectra of extracellular potentials from STN, VIM and GPi show no knee and lower aperiodic exponent than cortical recordings. A) Anatomical localizations of cortical and subcortical electrodes from the DBS lead, relative to the STN, GPi and VIM (Distal and Morel atlases, respectively). B) Representative example of power spectra, aperiodic component (gray lines) and model fit (dashed lines) for a STN and a cortical contact from the same subject. Note that for visual clarity the full-model fits were displaced vertically as indicated by the colored arrows on the left of the plot. C) Same as B for VIM. D) Same as B for GPi. E) Distribution of fitted aperiodic exponents for STN, VIM and GPi compared to cortex in individual subjects. Each dot corresponds to the median and error-bars to the standard deviation of all electrodes within the corresponding brain region. Gray lines join subcortical and cortical values for individual subjects. F) Same as E for the aperiodic knee frequency. Note that the y axis is in log-scale. The dashed horizontal line represents the smallest positive frequency acquired f_min_. Fits with knee frequencies smaller than f_min_ indicate spectra without observable knee. G) Same as E for the aperiodic offset.

The distribution of aperiodic parameters in STN recordings is remarkably different to that in cortical ECoG signals from the same subjects (Figure 4e, 4f and 4g, left panels). The aperiodic exponent for the STN has a median of 1.30±0.21 (median ± standard deviation across subjects), almost 3-fold smaller than that of ECoG recordings 3.41±0.30 for the same subjects (p<10^−5^, paired t-test, Table 1, Figure 4e). This difference in the aperiodic exponent between cortical and STN recordings reaches significance for all individual subjects analyzed (Figure 4e). Contrary to what we observed for cortical recordings, there was no correlation between the aperiodic slope from the STN and preoperative UPDRS-III ON or OFF scores (p=0.9 and p=0.7 respectively, Pearson correlation). The aperiodic exponent from DBS lead recordings in VIM and GPi were also significantly different from the simultaneous cortical recordings in each patient (p<10^−5^ for VIM, p=0.03 for GPi, paired t-test, Figure 4e).

**Table 1.** Mean and dispersion of aperiodic parameters across patient cohorts and brain structures. Exponent: Mean ± Standard deviation across patients. Offset (μV^2^/Hz): Median [Q_16_; Q_84_], note the asymmetric distribution. **f_*k*_**: Knee frequency in Hz, Median [Q_16_; Q_84_]. *P*(f_*k*_ < f_*min*_): percentage (±standard error) of electrodes with knee frequency lower than f_min_. *τ* = (2πf_*k*_)^−1^: aperiodic neural timescale in milliseconds. Abbreviations: PD, Parkinson’s disease; ET, essential tremor; EP, epilepsy; STN, subthalamic nucleus; VIM, ventral intermedius nucleus of the thalamus; GPi, globus pallidus internus; N_s_, number of subjects.

| Cohort | Location | Electrode type | N<sub>s</sub> | Exponent | Offset (μV<sup>2</sup>/Hz) | f<sub>k</sub> (Hz) | P(f<sub>k</sub> < f<sub>min</sub>) | τ (ms) |
| --- | --- | --- | --- | --- | --- | --- | --- | --- |
| PD<sub>STN-DBS</sub> | STN | DBS lead | 13 | 1.30±0.21 | 7.6 [2.9; 20] | < 1 | 85±4% | > 159 |
|  | Cortex | ECoG | 26 | 3.41±0.30 | 55 [15; 208] | 17.6 [13.3; 23.3] | 1.1±0.2% | 9.0 [6.8; 12.7] |
| ET<sub>VIM-DBS</sub> | VIM | DBS lead | 15 | 1.42±0.14 | 11.7 [6.0; 23] | < 1 | 87±3% | > 159 |
|  | Cortex | ECoG | 18 | 3.20±0.36 | 38 [11; 133] | 17.0 [12.9; 22.6] | 0.9±0.2% | 9.4 [7.0; 12.3] |
| PD<sub>GPI-DBS</sub> | GPI | DBS lead | 3 | 1.43±0.10 | 17.8 [8.4; 38] | < 1 | 73±9% | > 159 |
|  | Cortex | ECoG | 4 | 2.91±0.32 | 63 [42; 94] | 11.4 [7.2; 18.2] | 1.6±0.6% | 13.9 [8.7; 22.1] |
| EP<sub>sEEG</sub> | Thalamus | sEEG | 8 | 1.33±0.23 | 4.2 [1.0; 18.2] | < 1 | 79±4% | > 159 |
|  | Cortex | sEEG | 8 | 2.96±0.36 | 96 [14; 664] | 7.6 [3.1; 18.3] | 3.7±1.6% | 20.9 [8.7; 51.3] |

The calculated aperiodic knee frequency also exhibited a strikingly different distribution for STN, VIM and GPi than for the cortex (Table 1, Figure 4f). While the cortical knee frequencies center at 17±5 Hz (median ± standard deviation across subjects), those for STN, GPi and VIM converge to values lower than the smallest positive frequency of the power spectra (f_min_ = 1 *Hz*, dashed line in Figure 4f), in many cases reaching the lower boundary allowed for the fitting algorithm (0.1 Hz). It is important to note that knee frequency values smaller than f_min_ should not be interpreted quantitively, instead, they indicate the absence of a knee in the power spectra within the frequency ranged acquired. In other words, if there is a knee for the power spectra of STN, GPi and VIM, this value is lower than 1 Hz. Due to the high-pass frequency filters applied at acquisition it is not possible to explore lower frequencies in this dataset. The proportion of power spectra without a knee *P*(*f_k_* < *f_min_*) is significantly higher for STN, VIM and GPi recordings than for cortical recordings (Table 1, p<10^−6^, Fisher test).

Given the large difference observed in aperiodic parameters for STN, GPi and VIM as compared to cortex, we asked if these differences are specific to the types of electrodes used to record from subcortical nuclei in movement disorder patients or, on the contrary, generalize to other electrode types, subcortical structures, and diagnoses. To this end, we explored baseline recordings from 8 epilepsy patients undergoing intracranial monitoring with electrodes implanted in the thalamus for the purpose of assessing thalamic participation in the hypothesized seizure network and potential for therapeutic neuromodulation (Richardson 2022) (Figure 5a). In these recordings, the same type of stereo-EEG electrode contacts, and in some cases contacts on the same electrode, were used for cortical and thalamic targets. Thalamic contacts covered several thalamic nuclei from the ventral division (VLpd, VPLp, VLpv, VLa, VPM) to intralaminar nuclei (CM, MDpc, Pf, CL) (Morel, 2007) (see Table S3), whereas selected cortical contacts covered parietal and frontal regions (Figure 5a). As before, we found that thalamic power spectra show no observable knee, whereas cortical spectra from the same patients show prominent aperiodic knees (Figure 5b).

**Figure 5.**
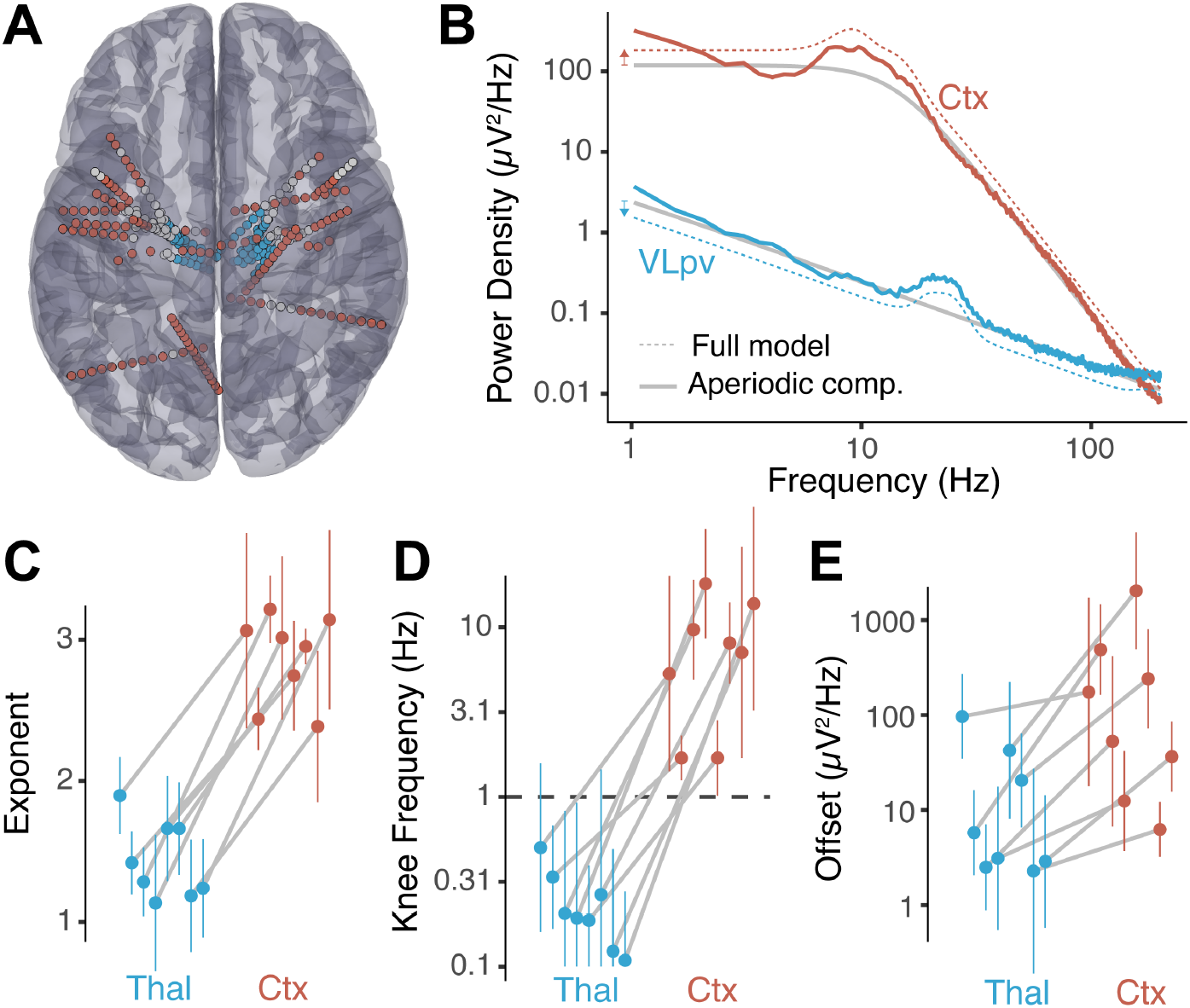
Power spectra from thalamic sEEG recordings show no knee and lower aperiodic exponent than cortical sEEG signals. A) Anatomical localizations of selected sEEG electrodes for epilepsy patients with thalamic implantations. B) Representative example of power spectra aperiodic component (gray line) and model fit (dashed lines) from a cortical sEEG contact (red) and thalamic sEEG (blue) recordings. For visual clarity, the full-model fits were displaced vertically as indicated by the colored arrows on the left. C) Distribution of fitted aperiodic exponents for thalamic bipolar pairs compared to cortex in individual subjects. Each dot corresponds to the median and the error-bars to the standard deviation of all bipolar pairs within the thalamus and cortex. The gray lines join parameters of the same subject. D) Same as C for the knee frequency. Note that the y axis is in log-scale. The dashed horizontal line represents f_min_. E) Same as C for the aperiodic offset.

A large difference in aperiodic exponent between cortical and thalamic electrodes was observed (p<0.001, paired t-test, Figure 5c, Table 1), consistent with the results obtained from movement disorder patients (Figure 4e). This difference holds at the single subject level, showing consistent changes across subjects (Figure 5c, gray lines). The aperiodic knee frequency also showed significant differences for thalamic and cortical contacts (Figure 5d), with thalamic values falling almost exclusively below f_min_ (smallest positive frequency of the spectra) and cortical values above this threshold (p<10^−6^, Fisher exact test).

Note that the aperiodic exponent of thalamic sEEG recordings in epilepsy patients (1.33±0.23) was not significantly different than that of DBS lead recordings in movement disorder patients (Table 1, p>0.05 for all pairwise comparisons by FDR-corrected permutation test). Similarly, the knee frequency extracted was below the cut-off value of f_min_=1Hz, as was the case for DBS recordings.

## Discussion

Almost every cortical region projects to and receives projections from the thalamus and other subcortical structures (Caviness and Frost, 1980; Sherman, 2016). These interactions provide a substrate for communication between distant cortical regions, facilitating spatial integration of the brain (Grant et al., 2012) and creating circuits with massive convergence and divergence in cell number at different nodes, as in the cortico-basal ganglia-thalamo-cortical loop (Bergman, 2021; Wilson, 2013). This organization involves regions whose cell types differ on many levels including channel and receptor expression, morphology, cytoarchitectures, and proportions of excitatory and inhibitory interactions, differences that allow for distinct dynamical behaviors and computational properties across brain structures.

In this study we performed systematic analysis of the broadband aperiodic component of brain recordings from multiple locations of the cortico-basal ganglia-thalamo-cortical loop, by fitting a phenomenological model to the power spectra of LFPs (Donoghue et al., 2020; Haller et al., 2018). We developed a novel parameterization of the broadband aperiodic component with the following advantages: 1) well-defined units for all parameters, 2) easily interpretable parameters, 3) structurally uncorrelated parameters, 4) parameters with more constrained physiological ranges, and 5) ability to fit spectra with or without an aperiodic ‘knee’ using the same model (see Figure 2a-b and Supplementary Materials). Interestingly, even with the novel parameterization of the aperiodic exponent, which removes structural correlations between parameters (Cedersund and Roll, 2009), residual positive correlation between the aperiodic knee and the exponent of the spectra (Figure S2b) was observed, suggesting that these parameters could be coupled.

Using this model to fit power spectra from baseline ECoG recordings from patients undergoing DBS implantation surgery, we found that the cortical aperiodic exponent correlates with Parkinson’s disease severity as assessed by the pre-operative UPDRS III (on-medication, Figure 3b). This novel result is in line with a MEG finding showing higher aperiodic exponents for PD patients compared to neuro-typical controls (Vinding et al., 2020). In our data, the correlation with the aperiodic exponent did not reach significance for the pre-operative UPDRS-OFF score, even though patients were in an OFF state during the intra-operative recordings. This could be due to less sensitivity or higher variability for the UPDRS-OFF score (as compared to the ON score) for the clinical population undergoing DBS treatment, which is biased to high symptom severity. There were no significant correlations of the STN LFP aperiodic exponent with UPDRS-III score (ON nor OFF levodopa), consistent with a recent report (Wiest et al., 2022).

Total beta power is known to correlate with PD disease severity in the basal ganglia (Brown et al., 2001; Cassidy et al., 2002; Kühn et al., 2004) and sensory-motor cortex (Pollok et al., 2012; Williams et al., 2002). FOOOF was designed to decouple oscillations from the underlying broadband aperiodic component, which reflects features of the entire spectrum, not just a specific band. However, estimations of aperiodic parameters can be affected by oscillatory components that extend beyond the fitting range (Gerster et al., 2022). This is not the case for our data since beta (12-30Hz) frequencies are above the lower frequency acquired (f_min_=1Hz). Additionally, we included a regularization term penalizing peaks below f_min_ to avoid this pitfall (see methods). Indeed, the fact that we obtained a significant correlation of the aperiodic exponent with UPDRS for motor cortex but not in the basal ganglia (which has prominent pathological beta oscillations), suggests that the method is correctly decoupling the aperiodic component from oscillatory features. Interestingly, changes in the aperiodic exponent could contribute to the known correlation of total beta power with UPDRS (Pollok et al., 2012; Williams et al., 2002) and to the ability of algorithms based on holistic spectral features to differentiate PD patients from controls (Anjum et al., 2020).

The main finding of this work is the conspicuous difference in the aperiodic component of the spectra between cortical recordings and those of basal ganglia and thalamic nuclei (Figure 4 and 5). Whereas cortical recordings showed an aperiodic knee with significant changes across cortical regions (Figure 3e, consistent with recent reports (Chaoul and Siegel, 2021; Gao et al., 2020b; Muthukumaraswamy and Liley, 2018)), spectra from basal ganglia and thalamic nuclei show no knee, an observation we could systematically evaluated thanks to the novel parameterization of the broadband aperiodic component. Spectra from subcortical regions showed an aperiodic exponent close to one (*χ* = 1.3 ± 0.2), significantly smaller than in cortex (*χ* = 3.2 ± 0.3). These results are reproducible across patients, two medical centers, electrode types, recording systems, referencing montages, diagnoses, and subcortical structures. Furthermore, the value for the aperiodic exponent in the STN we measured is consistent with recent studies that estimated this parameter (Huang et al., 2020; Wiest et al., 2022).

A limitation of this work is that ECoG electrodes lie over the pia mater, whereas the DBS leads penetrate the brain parenchyma. However, our data from epilepsy patients was recorded from the same type of sEEG electrodes for cortical and thalamic regions. Notably, these multi-contact electrodes are similar in size, shape, and impedance value to DBS lead contacts (Supplementary Table S1). We observed the same qualitative difference in aperiodic parameters between cortical and subcortical regions in both datasets, suggesting that these differences cannot be fully explained by electrode type and are due to structural and/or functional properties of the recorded brain areas. Another important limitation of our work is that different subcortical regions were recorded from different clinical populations. Therefore, we did not compare parameters across subcortical regions since the pathology would be an unavoidable confound; we limited our analysis of cortical vs. subcortical aperiodic activity to within-subject comparisons.

Neural morphology affects the shape and amplitude of extracellular potentials and could explain the differences in aperiodic activity observed between cortical and subcortical structures. Cells with large spatial separation between current sinks and return currents (like cortical pyramidal neurons) induce substantial extracellular ionic flows and large perturbations of the extracellular potential (Johnston and Wu, 1995). In contrast, neurons with roughly spherically symmetric dendritic arbors (like thalamocortical or STN neurons) do not produce strong current dipoles, with smaller contributions to recorded extracellular field potentials (Buzsáki et al., 2012; Johnston and Wu, 1995). However, synaptic inputs to subcortical structures may have asymmetric distributions which can produce measurable field potentials (Buzsáki et al., 2012; Lindén et al., 2010; Tanaka and Nakamura, 2019), for example having inhibitory synapses closer to the soma and more distal excitatory inputs (Lempka and McIntyre, 2013; Mazzoni et al., 2015; Wilson, 2010).

Although neuronal densities are comparable between cortical gray matter, STN and VIM (Bergman, 2021; Lévesque and Parent, 2005), the spatial arrangement of neurons can also have a large effect on the recorded extracellular potential (Gold et al., 2006; Johnston and Wu, 1995; Pettersen et al., 2008). In neuronal populations organized in layers, such as the 6-layered neocortex, simultaneous contributions from multiple similarly oriented cells will add up to give large fluctuations of the extracellular potential. In contrast, in neurons that have spatially isotropic arrangements, as in subcortical nuclei, simultaneous contributions from different units in diverse orientation can cancel out to some extent, producing overall smaller extracellular potentials (Johnston and Wu, 1995). These structural differences can explain why the overall power of field potentials is lower in subcortical nuclei than in neocortex. However, they do not explain why the aperiodic exponent and knee are different across these structures.

Several mechanisms have been suggested as the origin of the 1/*f^χ^* aperiodic component, including ionic diffusion and induction of electric fields in passive cells (Bédard et al., 2006a; Bédard and Destexhe, 2009). However, these effects are likely to be present in all brain regions. The shape and length of the dendrites, along with the location of the synaptic input can give rise to different frequency dependences of the intrinsic dendritic filtering (Lindén et al., 2010). Due to the different morphology of cortical versus thalamic and basal ganglia neurons, this could contribute to the difference in 1/*f^χ^* slope observed. However, the aperiodic slope in the STN has been shown to change with Propofol anesthesia (Huang et al., 2020), dopaminergic medication and DBS treatment (Wiest et al., 2022), demonstrating that this parameter depends on dynamical aspects of neural activity and cannot be fully explained by morphology and cytoarchitecture.

Functional differences like the profile and characteristic duration of post-synaptic currents can affect the aperiodic slope. For example, sharp rise and exponential decays for post-synaptic currents give rise to a 1/*f*^2^ decline of power (Bédard et al., 2006b; Miller et al., 2009; Milstein et al., 2009), and the ratio of excitatory to inhibitory inputs can affect the aperiodic knee frequency of the spectra (Gao et al., 2017). Transitions between UP and DOWN states (i.e., rapid trains of correlated synaptic inputs followed by quiescent periods), can give rise to power spectra following 1/*f*^2^ decline (Baranauskas et al., 2012; Milstein et al., 2009). In contrast, Poissonian inputs uncorrelated across cells do not contribute to the frequency dependency of the spectra (Bédard et al., 2006b; Miller et al., 2009; Milstein et al., 2009). Interestingly, there is a surprisingly low spike-timing correlation in the pallidum (Bar-Gad et al., 2003; Nini et al., 1995; Raz et al., 2000) and structures with strong pallidal input, including GPi, STN and several nuclei of the ventral thalamus will have low input correlation, which contribute to the low amplitude (Lindén et al., 2011) and slow decline with frequency of the power spectra in these regions.

There is currently no consensus on the physiological interpretation of the aperiodic knee and its change across brain structures. Miller et al. showed in ECoG recordings an aperiodic slope of χ=2 for frequencies above 15 Hz up to a “knee” around 75 Hz, at which the aperiodic slope changed to χ=4, implying the existence of a characteristic time scale *τ* = (2π*f_k_*)^−1^ = 2 – 4*ms* (Miller et al., 2009). Using similar reasoning on the knee observed around 10 Hz, Gao et al. proposed the existence of an “aperiodic neural timescale” (of around 10 to 50 ms) that can be interpreted as the characteristic duration of an aperiodic fluctuation of the LFP (Gao et al., 2020b). In our data, this timescale is in the range of 10 to 20 ms (Table 1) and changes across cortical locations (Figure 3e), which is consistent with previous findings and suggests that this parameter might be reflecting an intrinsic feature of cortical micro-circuitry and computation (Gao et al., 2020b).

The lack of an observable aperiodic knee for thalamic and basal ganglia recordings (i.e., the fitted value is lower than the cut-off frequency *f*_min_; Figure 4 and 5) can be interpreted as reflecting the absence of any characteristic duration of aperiodic fluctuations (strict 1/*f* power law). However, the neural morphology and cytoarchitecture of these regions might prevent characteristic aperiodic fluctuations from being reflected in LFPs. Alternatively, the aperiodic neural timescale could be longer than what can be detected by our method due to technical limitations. The latter interpretation puts a lower bound of 129ms for the subcortical aperiodic neural timescale (*τ* > (2*π f*_min_)^−1^ = 129*ms* for *f*_min_ =1 *Hz*) and suggests that basal ganglia and ventral thalamic nuclei are slower than cortex in terms of their aperiodic fluctuations. Although speculative, this interpretation suggests that the basal ganglia-thalamo-cortical loop could be a site of temporal-integration, a notion that aligns well with the known role of this circuit in spatial-integration, action selection and motor control (Bergman, 2021; DeLong and Wichmann, 2010; Grant et al., 2012; Mink, 1996; Turner and Desmurget, 2010).

## Acknowledgments

We would like to thank our patient-participants for their time and effort. This work was funded by the National Institute of Health (U01NS098969 and U01NS117836 to R.M.R.).

## Data availability

The data of this study is hosted in the Data Archive BRAIN Initiative (DABI, https://dabi.loni.usc.edu/dsi/1U01NS098969) and is available upon request.

## Author Contributions

W.J.L. and R.M.R. performed the intraoperative recordings. A.B., W.J.L., and V.K. reconstructed the electrode localizations. A.B., W.J.L., V.K., and J.Z. preprocessed electrophysiological data. A.B. and J.Z. developed and implemented the model and analyzed data. A.B. and R.M.R. conceived the study and wrote the manuscript. All authors discussed the results and commented on the manuscript. The authors declare no competing financial interests.

## References

Anjum MF, Dasgupta S, Mudumbai R, Singh A, Cavanagh JF, Narayanan NS (2020) Linear predictive coding distinguishes spectral EEG features of Parkinson’s disease. Parkinsonism Relat D 79:79–85.

Avants BB, Epstein CL, Grossman M, Gee JC (2008) Symmetric diffeomorphic image registration with cross-correlation: Evaluating automated labeling of elderly and neurodegenerative brain. Med Image Anal 12:26–41.

Baranauskas G, Maggiolini E, Vato A, Angotzi G, Bonfanti A, Zambra G, Spinelli A, Fadiga L (2012) Origins of 1/f 2 scaling in the power spectrum of intracortical local field potential. J Neurophysiol 107:984–994.

Bar-Gad I, Heimer G, Ritov Y, Bergman H (2003) Functional Correlations between Neighboring Neurons in the Primate Globus Pallidus Are Weak or Nonexistent. J Neurosci 23:4012–4016.

Basar E, Güntekin B (2013) Review of delta, theta, alpha, beta and gamma response oscillation in neuropsychiatric disorders In: Application of Brain Oscillations in Neuropsychiatric Diseases, Supplements to Clinical Neurophysiology (Basar E., Basar-Eroglu C, Ozerdem A, Rossini PM, Yener GG eds), pp303–341.

Bédard C, Destexhe A (2009) Macroscopic Models of Local Field Potentials and the Apparent 1/f Noise in Brain Activity. Biophys J 96:2589–2603.

Bédard C, Kröger H, Destexhe A (2006a) Model of low-pass filtering of local field potentials in brain tissue. Phys Rev E 73:051911.

Bédard C, Kröger H, Destexhe A (2006b) Does the 1/f Frequency Scaling of Brain Signals Reflect Self-Organized Critical States? Phys Rev Lett 97:118102.

Berger H (1929) Über das Elektrenkephalogramm des Menschen. Archiv Für Psychiatrie Und Nervenkrankheiten 87:527–570.

Bergman H (2021) The hidden life of the basal ganglia. Cambridge, Massachusetts: MIT Press.

Bretz F, Hothorn T, Westfall P (2011) Multiple Comparisons Using R. CRC Press.

Brown P, Oliviero A, Mazzone P, Insola A, Tonali P, Lazzaro VD (2001) Dopamine Dependency of Oscillations between Subthalamic Nucleus and Pallidum in Parkinson’s Disease. J Neurosci 21:1033–1038.

Bush A, Chrabaszcz A, Peterson V, Saravanan V, Dastolfo-Hromack C, Lipski WJ, Richardson RM (2021) Differentiation of speech-induced artifacts from physiological high gamma activity in intracranial recordings. Biorxiv 2021.04.26.441553.

Buzsáki G, Anastassiou CA, Koch C (2012) The origin of extracellular fields and currents — EEG, ECoG, LFP and spikes. Nat Rev Neurosci 13:407–420.

Cassidy M, Mazzone P, Oliviero A, Insola A, Tonali P, Lazzaro VD, Brown P (2002) Movement-related changes in synchronization in the human basal ganglia. Brain 125:1235–1246.

Caviness VS, Frost DO (1980) Tangential organization of thalamic projections to the neocortex in the mouse. J Comp Neurol 194:335–367.

Cedersund G, Roll J (2009) Systems biology: model based evaluation and comparison of potential explanations for given biological data. Febs J 276:903–922.

Chaoul AI, Siegel M (2021) Cortical correlation structure of aperiodic neuronal population activity. Neuroimage 245:118672.

Colombo MA, Napolitani M, Boly M, Gosseries O, Casarotto S, Rosanova M, Brichant J-F, Boveroux P, Rex S, Laureys S, Massimini M, Chieregato A, Sarasso S (2019) The spectral exponent of the resting EEG indexes the presence of consciousness during unresponsiveness induced by propofol, xenon, and ketamine. Neuroimage 189:631–644.

Dave S, Brothers TA, Swaab TY (2018) 1/f neural noise and electrophysiological indices of contextual prediction in aging. Brain Res 1691:34–43.

DeLong M, Wichmann T (2010) Changing Views of Basal Ganglia Circuits and Circuit Disorders. Clin Eeg Neurosci 41:61–67.

Donoghue T, Haller M, Peterson EJ, Varma P, Sebastian P, Gao R, Noto T, Lara AH, Wallis JD, Knight RT, Shestyuk A, Voytek B (2020) Parameterizing neural power spectra into periodic and aperiodic components. Nat Neurosci 23:1655–1665.

Engel AK, Fries P, Singer W (2001) Dynamic predictions: Oscillations and synchrony in top–down processing. Nat Rev Neurosci 2:704–716.

Ewert S, Plettig P, Li N, Chakravarty MM, Collins DL, Herrington TM, Kühn AA, Horn A (2018) Toward defining deep brain stimulation targets in MNI space: A subcortical atlas based on multimodal MRI, histology and structural connectivity. Neuroimage 170:271–282.

Fischl B, Salat DH, Busa E, Albert M, Dieterich M, Haselgrove C, Kouwe A van der, Killiany R, Kennedy D, Klaveness S, Montillo A, Makris N, Rosen B, Dale AM (2002) Whole Brain Segmentation Automated Labeling of Neuroanatomical Structures in the Human Brain. Neuron 33:341–355.

Fonov V, Evans AC, Botteron K, Almli CR, McKinstry RC, Collins DL, Group the BDC (2011) Unbiased average age-appropriate atlases for pediatric studies. Neuroimage 54:313–327.

Gao R, Brink RL van den, Pfeffer T, Voytek B (2020a) Neuronal timescales are functionally dynamic and shaped by cortical microarchitecture. Biorxiv 2020.05.25.115378.

Gao R, Brink RL van den, Pfeffer T, Voytek B (2020b) Neuronal timescales are functionally dynamic and shaped by cortical microarchitecture. Elife 9:e61277.

Gao R, Peterson EJ, Voytek B (2017) Inferring synaptic excitation/inhibition balance from field potentials. Neuroimage 158:70–78.

Gerster M, Waterstraat G, Litvak V, Lehnertz K, Schnitzler A, Florin E, Curio G, Nikulin V (2022) Separating Neural Oscillations from Aperiodic 1/f Activity: Challenges and Recommendations. Neuroinformatics 20:991–1012.

Glasser MF, Coalson TS, Robinson EC, Hacker CD, Harwell J, Yacoub E, Ugurbil K, Andersson J, Beckmann CF, Jenkinson M, Smith SM, Essen DCV (2016) A multi-modal parcellation of human cerebral cortex. Nature 536:171–178.

Gold C, Henze DA, Koch C, Buzsáki G (2006) On the Origin of the Extracellular Action Potential Waveform: A Modeling Study. J Neurophysiol 95:3113–3128.

Grant E, Hoerder-Suabedissen A, Molnár Z (2012) Development of the Corticothalamic Projections. Front Neurosci-switz 6:53.

Haller M, Donoghue T, Peterson E, Varma P, Sebastian P, Gao R, Noto T, Knight RT, Shestyuk A, Voytek B (2018) Parameterizing neural power spectra. Biorxiv 299859.

Horn A et al. (2019) Lead-DBS v2: Towards a comprehensive pipeline for deep brain stimulation imaging. Neuroimage 184:293–316.

Hothorn T, Hornik K, Wiel MA van de, Zeileis A (2008) Implementing a Class of Permutation Tests: The coin Package. Wiley Interdiscip Rev Comput Statistics 1:128–129.

Huang Y, Hu K, Green AL, Ma X, Gillies MJ, Wang S, Fitzgerald JJ, Pan Y, Martin S, Huang P, Zhan S, Li D, Tan H, Aziz TZ, Sun B (2020) Dynamic changes in rhythmic and arrhythmic neural signatures in the subthalamic nucleus induced by anaesthesia and tracheal intubation. Brit J Anaesth 125:67–76.

Johnston D, Wu SM-S (1995) Foundations of cellular neurophysiology. The MIT Press.

Kühn AA, Williams D, Kupsch A, Limousin P, Hariz M, Schneider G, Yarrow K, Brown P (2004) Event-related beta desynchronization in human subthalamic nucleus correlates with motor performance. Brain 127:735–746.

Kuznetsova A, Brockhoff PB, Christensen RHB (2017) lmerTest Package: Tests in Linear Mixed Effects Models. J Stat Softw 82.

Lempka SF, McIntyre CC (2013) Theoretical Analysis of the Local Field Potential in Deep Brain Stimulation Applications. Plos One 8:e59839.

Lendner JD, Helfrich RF, Mander BA, Romundstad L, Lin JJ, Walker MP, Larsson PG, Knight RT (2020) An electrophysiological marker of arousal level in humans. Elife 9:e55092.

Lévesque J, Parent A (2005) GABAergic interneurons in human subthalamic nucleus. Movement Disord 20:574–584.

Lindén H, Pettersen KH, Einevoll GT (2010) Intrinsic dendritic filtering gives low-pass power spectra of local field potentials. J Comput Neurosci 29:423–444.

Lindén H, Tetzlaff T, Potjans TC, Pettersen KH, Grün S, Diesmann M, Einevoll GT (2011) Modeling the Spatial Reach of the LFP. Neuron 72:859–872.

Mazzoni A, Lindén H, Cuntz H, Lansner A, Panzeri S, Einevoll GT (2015) Computing the Local Field Potential (LFP) from Integrate-and-Fire Network Models. Plos Comput Biol 11:e1004584.

Miller KJ, Sorensen LB, Ojemann JG, Nijs M den (2009) Power-Law Scaling in the Brain Surface Electric Potential. Plos Comput Biol 5:e1000609.

Milstein J, Mormann F, Fried I, Koch C (2009) Neuronal Shot Noise and Brownian 1/f2 Behavior in the Local Field Potential. Plos One 4:e4338.

Mink JW (1996) THE BASAL GANGLIA: FOCUSED SELECTION AND INHIBITION OF COMPETING MOTOR PROGRAMS. Prog Neurobiol 50:381–425.

Miskovic V, MacDonald KJ, Rhodes LJ, Cote KA (2019) Changes in EEG multiscale entropy and power-law frequency scaling during the human sleep cycle. Hum Brain Mapp 40:538–551.

Morel A (2007) Stereotactic Atlas of the Human Thalamus and Basal Ganglia. Informa Healthcare.

Muthukumaraswamy SD, Liley DTJ (2018) 1/f electrophysiological spectra in resting and drug-induced states can be explained by the dynamics of multiple oscillatory relaxation processes. Neuroimage 179:582–595.

Nini A, Feingold A, Slovin H, Bergman H (1995) Neurons in the globus pallidus do not show correlated activity in the normal monkey, but phase-locked oscillations appear in the MPTP model of parkinsonism. J Neurophysiol 74:1800–1805.

Nunez PL, Srinivasan R (2006) Electric Fields of the Brain. Oxford University Press.

Oostenveld R, Fries P, Maris E, Schoffelen J-M (2011) FieldTrip: Open Source Software for Advanced Analysis of MEG, EEG, and Invasive Electrophysiological Data. Comput Intel Neurosc 2011:156869.

Pettersen KH, Hagen E, Einevoll GT (2008) Estimation of population firing rates and current source densities from laminar electrode recordings. J Comput Neurosci 24:291–313.

Pollok B, Krause V, Martsch W, Wach C, Schnitzler A, Südmeyer M (2012) Motor-cortical oscillations in early stages of Parkinson’s disease. J Physiology 590:3203–3212.

Randazzo MJ, Kondylis ED, Alhourani A, Wozny TA, Lipski WJ, Crammond DJ, Richardson RM (2016) Three-dimensional localization of cortical electrodes in deep brain stimulation surgery from intraoperative fluoroscopy. Neuroimage 125:515–521.

Raz A, Vaadia E, Bergman H (2000) Firing Patterns and Correlations of Spontaneous Discharge of Pallidal Neurons in the Normal and the Tremulous 1-Methyl-4-Phenyl-1,2,3,6-Tetrahydropyridine Vervet Model of Parkinsonism. J Neurosci 20:8559–8571.

Richardson RM (n.d.) Closed-Loop Brain Stimulation and Paradigm Shifts in Epilepsy Surgery. Neurol Clin 40:355–373.

Sherman SM (2016) Thalamus plays a central role in ongoing cortical functioning. Nat Neurosci 19:533–541.

Tadel F, Baillet S, Mosher JC, Pantazis D, Leahy RM (2011) Brainstorm: A User-Friendly Application for MEG/EEG Analysis. Comput Intel Neurosc 2011:879716.

Tanaka T, Nakamura KC (2019) Focal inputs are a potential origin of local field potential (LFP) in the brain regions without laminar structure. Plos One 14:e0226028.

Turner RS, Desmurget M (2010) Basal ganglia contributions to motor control: a vigorous tutor. Curr Opin Neurobiol 20:704–716.

Vinding MC, Tsitsi P, Waldthaler J, Oostenveld R, Ingvar M, Svenningsson P, Lundqvist D (2020) Reduction of spontaneous cortical beta bursts in Parkinson’s disease is linked to symptom severity. Brain Commun 2:fcaa052.

Voytek B, Kramer MA, Case J, Lepage KQ, Tempesta ZR, Knight RT, Gazzaley A (2015) Age-Related Changes in 1/f Neural Electrophysiological Noise. J Neurosci 35:13257–13265.

Welch PD (1967) The use of fast Fourier transform for the estimation of power spectra: a method based on time averaging overt short, modified periodigrams. IEEE transactions on audio and electroacoustics AU-15:70–73.

Wen H, Liu Z (2016) Separating Fractal and Oscillatory Components in the Power Spectrum of Neurophysiological Signal. Brain Topogr 29:13–26.

Wiest C, Torrecillos F, Pogosyan A, Bange M, Muthuraman M, Groppa S, Hulse N, Hasegawa H, Ashkan K, Baig F, Morgante F, Pereira EA, Mallet N, Magill PJ, Brown P, Sharott A, Tan H (2022) The aperiodic exponent of subthalamic field potentials reflects excitation/inhibition balance in Parkinsonism: a cross-species study in vivo. Biorxiv 2022.08.23.504923.

Williams D, Tijssen M, Bruggen G van, Bosch A, Insola A, Lazzaro VD, Mazzone P, Oliviero A, Quartarone A, Speelman H, Brown P (2002) Dopamine-dependent changes in the functional connectivity between basal ganglia and cerebral cortex in humans. Brain 125:1558–1569.

Wilson CJ (2013) Active decorrelation in the basal ganglia. Neuroscience 250:467–482.

Wilson CJ (2010) Subthalamo-Pallidal Circuit In: Handbook of Brain Microciircuits (Shepherd GM, Grillner S eds), pp127–134. Oxford University Press.

