## Supplementary Material for "Broadband aperiodic components of local field potentials reflect inherent differences between cortical and subcortical activity"

### Electrodes and acquisition systems

| TYPE | MODEL | DETAILS | SYSTEM | RATE (KHZ) | CH # | IMPEDANCE (KΩ) |
|---|---|---|---|---|---|---|
| DBS LEAD | Medtronic[1] 3389 | Pt/Ir rings. Diameter=1.27mm, length=1.5 mm, axial spacing=0.5 mm. | Grapevine[5] | 30 | 4 | 0.5 - 1.5 |
| DBS LEAD | St. Jude[2] 6172 short | Pt/Ir directional lead, 2 central rings split in 3. Diameter 1.27 mm, length=1.5 mm, axial spacing=0.5 mm. | Grapevine[5] | 30 | 8 | Ring: 0.5 - 1.5<br>Direct.: 1.0-3.0 |
| DBS LEAD | Boston Scientific[3] DB-2201-45 | Pt/Ir rings. Diameter 1.3 mm, length=1.5 mm, axial spacing=0.5 mm, contact span=15.5 mm. | Grapevine[5] | 30 | 8 | 0.5 - 1.5 |
| DBS LEAD | Boston Scientific[3] DB-2202-45 | Pt/Ir directional lead, 2 central rings split in 3. Diameter 1.3 mm, length=1.5 mm, axial spacing=0.5 mm, total span 7.5 mm. | Grapevine[5] | 30 | 8 | Ring: 0.5 - 1.5<br>Direct.: 1.0-3.0 |
| ECOG | PMT[4] Cortac 2110-54-001 | Pt. 1 mm disc contacts, 3x18 layout, 3 mm center-to-center | Grapevine[5] | 30 | 54 | 2.5 – 8.0 |
| ECOG | PMT[4] Cortac 2011-63-002 | Pt. 1 mm disc contacts, 3x63 layout, 3 mm center-to-center | Grapevine[5] | 30 | 63 | 2.5 – 8.0 |
| SEEG | PMT[4] sEEG Depth 2102-16 | Pt rings. Diameter=0.8mm, length=2m, axial spacing 3.5 to 4.43 mm (+ spacers) | Natus[6] | 1 | 16 | 1.2 - 2.1 [7] |
| SEEG | PMT[4] sEEG Depth 2102-08 | Pt rings. Diameter=0.8mm, length=2m, axial spacing 3.5 to 4.43 mm (+ spacers) | Natus[6] | 1 | 8 | 1.2 - 2.1 [7] |

**Supplementary Table S1.** Electrodes and acquisition system specifications for neural and physiological signals. [1] Medtronic, Minneapolis, MN, USA. [2] Abbott Neuromodulation, Austin, TX, USA. [3] Boston Scientific Neuromodulation Corp, Valencia, CA, USA. [4] PMT Corporation, Chanhassen, MN, USA. [5] Ripple LLC, Salt Lake City, UT, USA. [6] Natus Medical Incorporated, Pleasanton, CA. [7] As measured in (Carvallo et al., 2019).

### Lorentzian-like parameterization of the aperiodic component

In the original parameterization of the FOOOF model (Haller et al., 2018) the broadband aperiodic component of the power spectra was defined in log-power as $\log\left(L(f)\right) = \mathrm{b} - \log\left(\mathrm{k} + f^{\chi}\right)$, similar to the one proposed by (Miller et al., 2009). For an easier comparison with the novel parameterization proposed in this work we rewrite this expression in linear-power and add the subscript "o" for original.

$$\mathrm{L_o}(f) = \frac{\mathrm{A_o}}{\mathrm{k_o} + f^{\chi_o}}$$

where $b_o = \log(A_o)$.

Note that some parameters in this expression have ill-defined units. $L_o$ has units of power density $\mu V^2/Hz$, $f$ is a frequency and therefore has units of $Hz$, and $\chi_o$ is an adimensional exponent. Therefore, the parameter $k_o$ will have units of $Hz^{\chi_o}$ and $A_o$ will have units of $\mu V^2 Hz^{\chi_o - 1}$ (e.g., $A_o$ will have units of $\mu V^2$ if $\chi_o = 1$, but of $\mu V^2 Hz$ if $\chi_o = 2$). This means that the units of $k_o$ and $A_o$ depend on the value of $\chi_o$ and therefore are not properly defined, thus hindering the interpretation of these parameters.

We sought to correct this problem by proposing a new Lorentzian-like parameterization of the broadband aperiodic component with parameters with well-defined units.

$$L(f) = A\frac{f_k{}^{\chi} + f_{min}^{\chi}}{f_k{}^{\chi} + f^{\chi}}$$

Note that in this case $f_k$ has units of frequency (Hz), and A has units of power density ($\mu V^2/Hz$), independently of the value of $\chi$. $f_{min}$ is the smallest frequency for which a power estimation is available, which will be ultimately limited by the filtering characteristics of the data acquisition system. Note that this frequency will necessary be positive ($f_{min} > 0$) since virtually all amplifiers are AC coupled and therefore filter out the DC component. Note that $f_{min}$ is known *a priori* based on acquisition and/or analysis parameters and it is not a fitted during the optimization.

There is a simple mapping between the original and current parameters given by

$$\chi = \chi_o$$
$$f_k = k_o{}^{1/\chi_o}$$
$$A = A_o/(k_o + f_{min}^{\chi_o})$$

Since there is a one-to-one mapping between the two parameterizations, both will converge to the same predictions for the aperiodic component of the spectra. However, we argue that the current parameterization has more easily interpretable parameters.

The predicted power of the current parameterization of the broadband aperiodic component at the smallest positive frequency will always be $L(f_{min}) = A$, therefore A can be directly interpreted as the aperiodic power at $f_{min}$ (Figure S1a). For the original parameterization the value at $f_{min}$ is given by $L_o(f_{min}) = A_o/(k_o + f_{min}^{\chi_o})$, which is linear with $A_o$, but also depends on other parameters. Therefore, the value of $A_o$ cannot be easily interpreted or read out from the y-axis of the plot (Figure S1b). More importantly, changing the value of $k_o$ can also affect the predicted value at $f_{min}$ (Figure S1b).

In the novel model, for cases in which $f_k \gg f_{min}$, the knee frequency can be interpreted as the frequency at which the predicted aperiodic power drops to half of that at $f_{min}$, since in this regime $L(f_k) = A/2 = L(f_{min})/2$ (Figure S1a). This simple interpretation is not valid for the original parameterization, as changing $k_o$ also changes the predicted power at $f_{min}$.

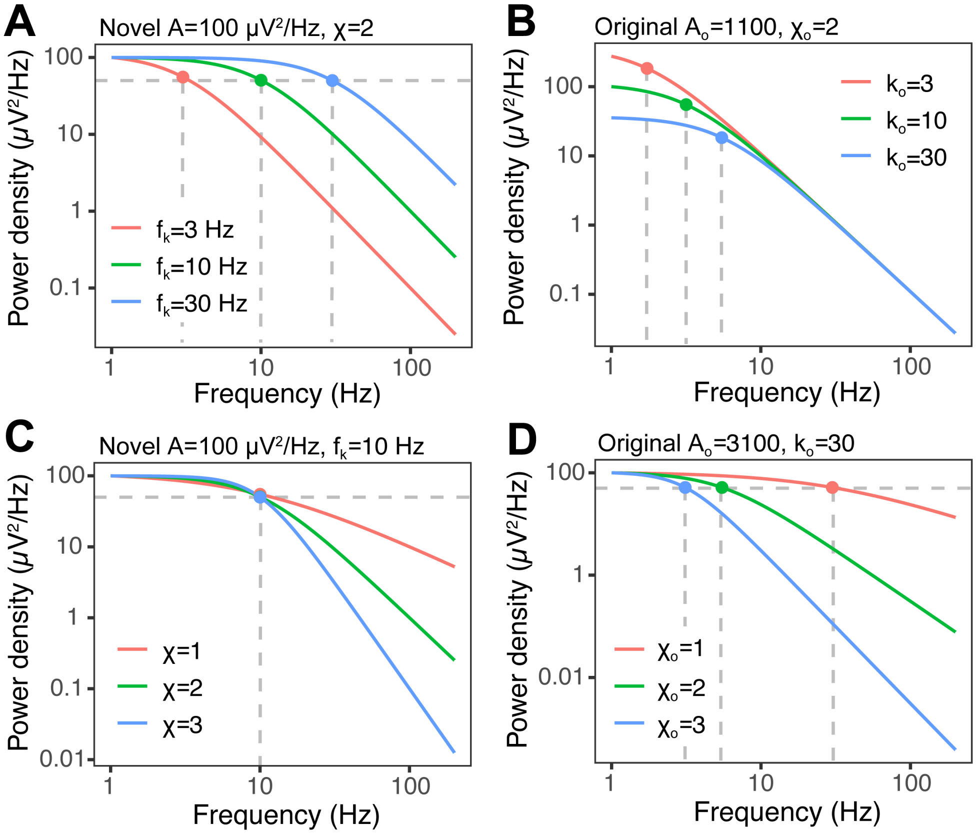


**Supplementary Figure S1:** Dependency of predicted aperiodic power density with parameters according to the novel (A and C) and original (B and D) parameterizations. A) Novel parametrization for three different knee frequencies (3 Hz, red curve; 10 Hz, green curve; 30 Hz, blue curve) with fixed values for A and $\chi$ indicated above the panel. The horizontal gray dashed line represents 50% of the power at $f_{min} = 1\ Hz$. Color dots and vertical dashed line represent the knee frequency for each curve. B) Same as A for the original parametrization for indicated values of $k_o$. C) Same as A for indicated values of $\chi$. D) Same as A for the original parametrization and indicated values of $\chi$.

Finally, the aperiodic exponents $\chi$ and $\chi_o$ are equivalent across parameterizations. However, changing this parameter only affects the slope of the decay with frequency in the novel parameterization (Figure S1c), whereas in the original model it can also change the frequency at which the power drops to 50% of the original value (Figure S1d).

Another advantage of the novel parameterization is that having $f_{min}$ explicitly defined allows to differentiate between regimes with and without a knee; $f_k \leq f_{min}$ should be interpreted as an aperiodic component without a detectable knee in the analyzed frequency range. In the original parameterization these two regimes were described by different models, referred to as "fixed" and "knee" models (Haller et al., 2018). Since $f_k$ is a positive scale parameter we decided to scan it in log-space during the fitting procedure, ensuring that it remains positive.

### Parameters correlation for original and current parametrizations

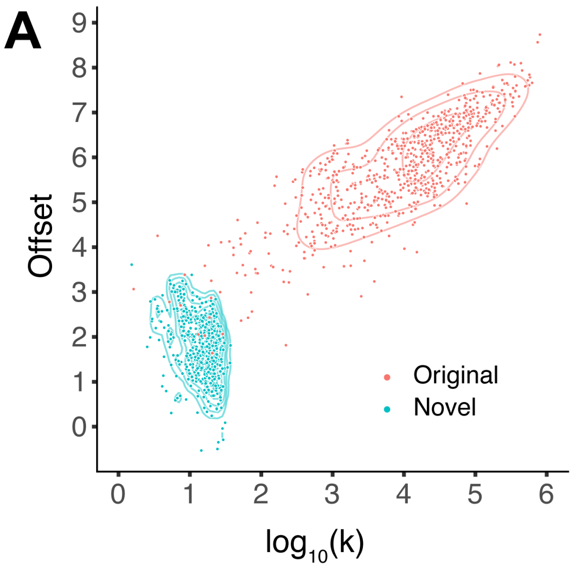


**Supplementary Figure S2:** A) Aperiodic offset vs. knee parameters for ECoG recordings from PD patients, for the original (red) and novel (blue) parameterizations. Contour lines represent the 5%, 10%, 20%, 40% and 80% percentiles of 2D kernel density estimation. B) Same as A for the aperiodic exponent vs. knee parameters.

### Aperiodic exponent versus UPDRS-III OFF and ON/OFF change percentage

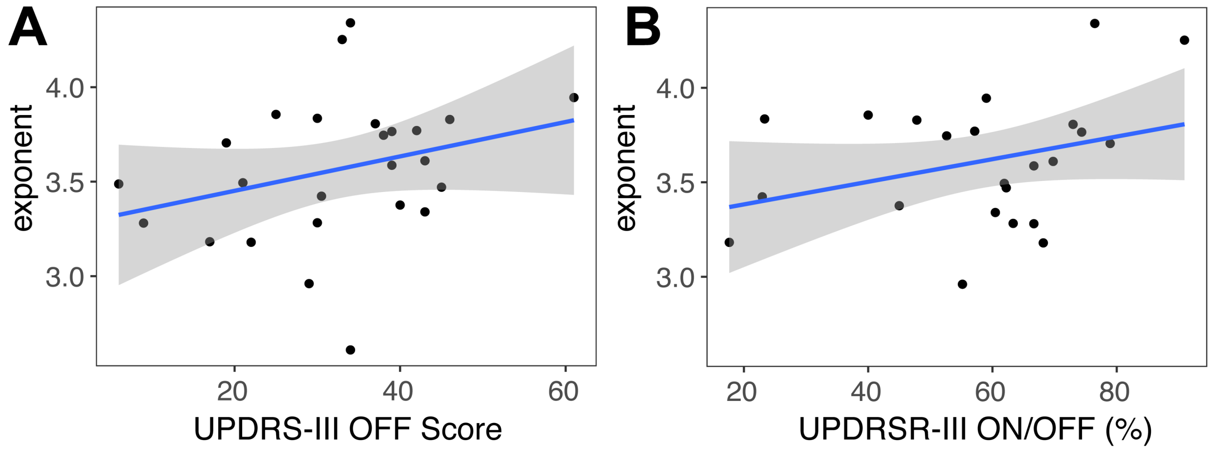


**Supplementary Figure S3:** A) Median cortical aperiodic exponent from Rolandic and premotor ECoG recordings in PD patients undergoing STN-DBS surgery vs. preoperative UPDRS-III score OFF dopaminergic medication. Shaded region represents CI95.B) Same as A but correlation against the ON/OFF percental change of preoperative UPDRS-III score.

### Aperiodic parameters by cortical region

| Label | $N_s$ | $f_k\ (Hz)$ | $\chi$ | $A\ \left(\frac{\mu V^2}{Hz}\right)$ | $Log_{10}(f_k)$ Region effect | $\chi$ Region effect | $Log_{10}(A)$ Region effect |
|---|---|---|---|---|---|---|---|
| *46v* | 12 | 16.5±1.5 | 3.49±0.10 | 116±33 | -0.006±0.036 | 0.112±0.090 | 0.228±0.051 |
| *8C* | 25 | 17.5±1.0 | 3.44±0.09 | 68±19 | -0.009±0.021 | 0.086±0.058 | 0.115±0.056 |
| *8Av* | 22 | 16.7±0.9 | 3.32±0.08 | 56±17 | -0.031±0.014 | -0.018±0.038 | 0.130±0.062 |
| *IFSp* | 13 | 17.7±1.2 | 3.39±0.11 | 55±22 | -0.007±0.021 | 0.011±0.073 | 0.013±0.084 |
| *44* | 18 | 17.1±0.9 | 3.20±0.09 | 28±8 | 0.000±0.017 | -0.104±0.063 | -0.201±0.084 |
| *6r* | 18 | 16.0±0.9 | 3.15±0.09 | 25±5 | -0.034±0.018 | -0.085±0.045 | -0.153±0.083 |
| *6v* | 41 | 18.1±0.6 | 3.48±0.08 | 57±13 | 0.001±0.009 | 0.153±0.055 | 0.146±0.040 |
| *55b* | 27 | 18.6±0.8 | 3.51±0.08 | 69±20 | 0.012±0.010 | 0.166±0.042 | 0.176±0.050 |
| *4* | 17 | 19.2±1.2 | 3.44±0.12 | 42±18 | -0.006±0.027 | 0.062±0.069 | 0.071±0.085 |
| *3a* | 14 | 18.1±1.2 | 3.24±0.16 | 20±8 | -0.019±0.021 | -0.089±0.089 | -0.248±0.071 |
| *3b* | 24 | 19.2±1.1 | 3.52±0.11 | 53±18 | 0.028±0.018 | 0.148±0.070 | 0.059±0.075 |
| *1* | 44 | 19.7±0.7 | 3.43±0.06 | 42±9 | 0.044±0.010 | 0.099±0.041 | -0.041±0.042 |
| *43* | 19 | 18.5±1.2 | 3.28±0.10 | 27±8 | -0.004±0.020 | -0.022±0.064 | -0.012±0.079 |
| *OP4* | 33 | 17.0±0.7 | 3.20±0.08 | 28±7 | -0.025±0.011 | -0.130±0.049 | -0.185±0.042 |
| *PFop* | 28 | 17.9±0.5 | 3.29±0.06 | 42±10 | -0.015±0.010 | -0.064±0.037 | -0.030±0.043 |
| *PF* | 18 | 18.4±1.0 | 3.30±0.06 | 46±13 | -0.002±0.010 | -0.055±0.051 | -0.028±0.065 |
| *PSL* | 9 | 16.9±1.2 | 3.23±0.10 | 71±28 | -0.040±0.023 | -0.186±0.073 | 0.016±0.025 |
| *STV* | 11 | 19.0±0.9 | 3.35±0.05 | 50±14 | -0.016±0.014 | -0.073±0.052 | 0.011±0.082 |
| *A5* | 24 | 17.1±0.8 | 3.24±0.07 | 36±9 | -0.033±0.016 | -0.079±0.041 | 0.040±0.068 |
| *A4* | 36 | 17.9±0.6 | 3.24±0.06 | 35±7 | -0.008±0.006 | -0.087±0.028 | -0.054±0.032 |

**Supplementary Table S2. Aperiodic parameters by cortical region. Label**: cortical label according to the multimodal parcellation 1 atlas (Glasser et al., 2016). $N_s$: number of subjects with electrodes in the region of interest. Only regions with electrodes from 9 or more subjects were included in the analysis. $f_k$**:** mean aperiodic knee frequency in Hz ± standard error of the mean across subjects (SE). $\chi$: mean aperiodic exponent ± SE. $A$: mean offset in $\mu V^2/Hz$ ± SE. $Log_{10}(f_k)$ **region effect:** logarithm of the aperiodic knee frequency after subtracting the subject effect ± SE. $\chi$ **region effect:** aperiodic exponent after subtracting the subject effect ± SE. $Log_{10}(A)$ **region effect:** logarithm of the aperiodic offset after subtracting the subject effect ± SE.

### Thalamic nuclei recorded with sEEG electrodes

| *Label* | $N_{subjects}$ | $N_{contacts}$ | **Full name (Morel atlas)** |
|---|---|---|---|
| *VLpd* | 6 | 17 | Ventral lateral posterior nucleus (dorsal division) |
| *CM* | 5 | 12 | Centromedian |
| *VPLp* | 5 | 7 | Ventral posterior lateral nucleus (posterior division) |
| *VLpv* | 4 | 11 | Ventral lateral posterior nucleus (ventral divisions) |
| *MDpc* | 4 | 9 | Mediodorsal nucleus (parvocellular division) |
| *Pf* | 4 | 6 | Parafascicular nucleus |
| *CL* | 3 | 6 | Central lateral nucleus |
| *VLa* | 3 | 6 | Ventral lateral anterior nucleus |
| *VPM* | 3 | 4 | Ventral posterior medial nucleus |
| *Hb* | 3 | 3 | Habenular nucleus |
| *LP* | 1 | 4 | Lateral posterior nucleus |
| *PuM* | 1 | 2 | Medial pulvinar |
| *Li* | 1 | 1 | Limitans nucleus |
| *PuA* | 1 | 1 | Anterior pulvinar |
| *VApc* | 1 | 1 | Ventral anterior nucleus (parvocellular divisions) |
| *VPLa* | 1 | 1 | Ventral posterior lateral nucleus (anterior divisions) |

**Supplementary Table S3. Coverage of thalamic nuclei in sEEG dataset.**

### References


**Carvallo A, Modolo J, Benquet P, Lagarde S, Bartolomei F, Wendling F (2019) Biophysical Modeling for Brain Tissue Conductivity Estimation Using SEEG Electrodes. IEEE TRANSACTIONS ON BIOMEDICAL ENGINEERING 66:1695–1704.**

**Glasser MF, Coalson TS, Robinson EC, Hacker CD, Harwell J, Yacoub E, Ugurbil K, Andersson J, Beckmann CF, Jenkinson M, Smith SM, Essen DCV (2016) A multi-modal parcellation of human cerebral cortex. Nature 536:171–178.**

**Haller M, Donoghue T, Peterson E, Varma P, Sebastian P, Gao R, Noto T, Knight RT, Shestyuk A, Voytek B (2018) Parameterizing neural power spectra. Biorxiv 299859.**

**Miller KJ, Sorensen LB, Ojemann JG, Nijs M den (2009) Power-Law Scaling in the Brain Surface Electric Potential. Plos Comput Biol 5:e1000609.**
